# Binder and monomer valencies determine the extent of collapse and reswelling of chromatin

**DOI:** 10.1101/2024.08.30.610569

**Authors:** Sougata Guha

## Abstract

Multivalent DNA-bridging proteins mediated collapse of chromatin polymer has long been established as one of the driving factors in chromatin organisation inside cells. These multivalent proteins can bind to distant binding sites along the chromatin backbone and bring them together in spatial proximity, leading to collapsed conformations. Recently, it has been suggested that these proteins not only drive collapse of the chromatin polymer, but also reswelling at higher concentrations. In this study, we investigate the physical mechanisms underlying this unexpected reswelling behaviour. We use Langevin dynamics simulation of a coarse-grained homopolymer to investigate the effects of the valencies of both the binders and the monomers on the polymer conformations. We find that while the extent of collapse of the polymer is strongly dependent on the binder valency, the extent of reswelling is largely determined by the monomer valency. Furthermore, we also discover two different physical mechanisms that drive reswelling of the polymer - *excluded volume effects* and *loss of long-range loops*. Finally, we obtain a classification map to determine the regimes in which each of these mechanisms is the dominant factor leading to polymer reswelling.

## I. INTRODUCTION

Understanding the physical principles behind the spatial and temporal organisation of chromatin inside the nucleus remains a central challenge. Dynamic chromatin organisation is critical for proper gene regulation, and perturbations in this assembly can lead to various diseases^1–6^. Recent developments in experimental techniques such as 3C, 4C, 5C, Hi-C, and micro-C have provided deep insight into the importance of chromosome contacts in chromatin organisation^7–14^. These experiments have revealed the hierarchical organisation of chromatin at multiple scales, through the formation of Topologically Associated Domains (TADs) and A/B compartments^15^. At larger scales, each chromosome occupies a distinct region within the nucleus, known as a chromosome territory, constrained by neighbouring chromosomes^15^.

Formation of chromatin loops is one of the ubiquitous structural motifs across cell types, and these loops are formed by different chromatin-bridging proteins^16–18^. These proteins bind to different segments of chromatin that are separated by a long distance along the chromatin chain and bring them spatially close to each other^19–24^. Long loops, driven by the Structural Maintenance of Chromatin (SMC) proteins such as cohesin and condensin, localise to TAD boundaries, and ensure insulation among neighbouring TADs. Chromatin organisation driven by loops is known to be spatially and temporally dynamic^25–27^, varying based on several factors such as chromatin epigenetic states^28,29^ and cell cycle phases^30,31^. Chromatin loops are also crucial in regulating various biological processes. Specific loops like TAD boundaries have been implicated in facilitating enhancer-promoter interactions^32–34^ which are necessary for gene activation and DNA replication^20,35^.

A critical factor underlying the formation of these loops is the interaction of the chromatin with various DNA-bridging proteins. These proteins are often multivalent i.e. they can bind to different numbers of chromatin segments simultaneously. For example cohesin and condensin crosslink chromatin segments and bring them spatially close to each other^7,32,36–38^. HP1 proteins are known to interact with methylated histone H3^39,40^ and are also reported to vary their valency through self oligomerisation^41,42^. Also, polycomb proteins (PRC1) are known to bring multiple DNA segments containing PRC1-binding domains together^43^. Patchy particles with limited valency have also been studied in both simulations^44,45^ and experiments^46,47^, as well as in the context of chromatin organisation^48^. Additionally, chromatin can also exhibit multivalency to many proteins. For example, PRC complexes bind to H2A or H3 histone proteins^49^. As each nucleosome consists of a pair of these histone proteins, nucleosomes themselves are bivalent to polycomb protein complexes. Also, in zebrafish sperm, developmental genes are packed in large blocks of chromatin that contain multiple binding sites of post-translational modification complexes^50^. Moreover, in a coarse-grained polymer picture of chromatin, each monomer consists of a chromatin segment that varies from nucleosome level (∼ 200*bp*)^51,52^ to 100 kbp^53^. These segments usually contain multiple binding domains for different proteins, and the number of such domains (i.e, chromatin valency) depends on the specific chromatin segment and the scale of coarse-graining. Therefore, understanding the impact of varying protein and chromatin valencies is critical in comprehending the processes of chromatin compaction and reswelling.

Polymer-based models of chromatin have yielded numerous insights into this process of protein-chromatin interactions and their role in regulating chromosome organisation^53–72^. A foundational model is the ‘Strings and Binders Switch’ (SBS) model^73,74^. In this model, chromatin is assumed to be a self-avoiding polymer, which consists of binding sites for diffusible bridging proteins. Diffusible bridging proteins dynamically attach to distant segments of the chromatin polymer and hence form loops. Due to this long-range bridging of chromatin segments, the chromatin polymer collapses and undergoes a coil-globule transition^56^. The transition point is known to be a function of binder concentration and chromatin-binder affinity^53^. The contact probability exponent and the polymer size exponents obtained in this model match very well with the experimentally observed exponents^75^. Subsequently, several variants of this model have been implemented in the context of chromatin organisation inside the cell nucleus. For example, incorporating the active and inactive epigenetic states of chromatin along with the semi-flexibility of chromatin leads to different collapsed structures of the chromatin clusters^57–59^. The presence of specific binding domains for different binders has been used to reproduce TADs and sub-TADs observed experimentally^67,76^. Many studies have also used binder-mediated bridging implicitly by considering intra-polymer crosslinking and explained various features of chromatin organisation^61,62,77–82^.

In a previous study^83^, we investigated the role of diffusible multivalent binders in chromatin organisation in confined systems. We reported that chromatin-protein interactions not only can drive polymer compaction, but also can reswell the polymer at higher concentrations. Additionally, the amount of reswelling, the required binder concentration for reswelling and the timescales are found to depend on the scale of confinement. This counterintuitive reswelling of the polymer has been studied extensively in the context of polymer cononsolvency where the polymer undergoes an open-collapsed-open transition in a mixture of two different solvents^84,85^. Interestingly, these studies were performed for relatively small polymer lengths (*N* ≤ 100) and at very high solvent number densities (*c* ≳ 0.7). Few recent studies have also reported this non-monotonic variation in polymer size in the context of chromatin organisation^86–88^. A critical factor in understanding this collapse and subsequent reswelling of chromatin is the interplay of binder and monomer valencies which have been ignored by all previous studies. These specific valencies of binders and chromatin determine the underlying looping kinetics, and therefore determine the structural properties of the chromatin polymer. In this work, we systematically study the effect of both the valency of the bridging protein and the monomer itself on the polymer conformations and spatial organisation.

## II. MODEL

To investigate how the valencies of both monomers and binders affect the collapse and reswelling of chromatin, we simulate a polymer-binder system immersed in an implicit good solvent under confinement. Using underdamped Langevin dynamics simulations, we model a polymer consisting of 512 spherical monomers, each with a diameter of *σ*. The polymer chain connectivity is maintained by introducing FENE potential^89^ between consecutive monomers and is given by

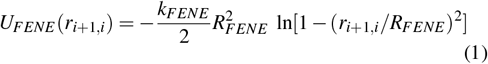

where *k*_*FENE*_ = 30*k*_*B*_*T* ^89^ is the spring constant, *R* = 1.6*σ* ^89^ is the maximum allowed bond length and *r*_*i, j*_ = |*r*_*i*_ − *r* _*j*_| is the euclidean distance between the *i*^*th*^ and *j*^*th*^ monomers. All non-bonded monomers interact via volume exclusion modelled as purely repulsive WCA potential^90^,

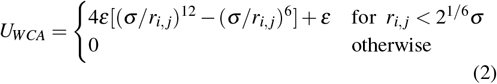

The interaction strength *ε* is chosen to be equal to *k*_*B*_*T*. Individual binders are modelled as spherical beads of diameter *σ*_*b*_ = *σ*. While the binders mutually interact via excluded volume interaction implemented by WCA potential (Eqn. 2), the polymer-binder interaction is modelled as ‘Truncated and Shifted Lennard-Jones’ (TSLJ) potential,

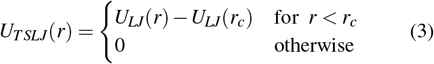

where *U*_*LJ*_(*r*) = 4*ε*_*m*_[(*σ/r*)^12^ ™ (*σ/r*)^6^] with *ε*_*m*_ = 3.5*ε*^83^ and *r*_*c*_ = 1.5*σ* ^83^. This interaction is permitted as long as both the monomer and the binder have not saturated their valency i.e., the monomer is bound to a number of binders and the binder is bound to a number of monomers less than their own valencies. If either the monomer or the binder has already saturated its valency, then they interact via volume exclusion. To ensure that the bonding between the monomers and the binders follows the valency restriction, we first make a ‘probable list’ of monomers that are within the interaction cut-off distance (*r*_*c*_) from a binder. Now, a number of monomers equal to the difference between binder valency and number of monomers bound to that binder is chosen at random from that list. The monomer valency restriction is also maintained while choosing the monomers. We repeat this method for all the binders at every given time step.

We implement the confinement as a spherical wall of radius *R* which interacts with all particles via WCA potential. In this paper, the chromatin volume density and confinement radius are kept fixed. We chose a confinement radius that corresponds to a volume density of chromatin, *φ* ∼ 0.044 (*R* ∼ 11.3*σ*), which is consistent with the density of chromatin observed experimentally within the nucleus, which varies in the range *φ* = 0.01 ™ 0.25^91^. We also keep the binder number density (*c*) within biologically relevant regime (*c* ∼ 0.04 ™ 0.3)^83,92^. Here, we investigate the conformations of chromatin polymers at different binder and monomer valencies. We discard the monovalency of the binders because, in such a case, formation of loops would not be possible, which would lead to no compaction of the polymer.

All simulations are performed using underdamped Langevin dynamics and integrations are performed using Velocity-Verlet algorithm^93^. There are two timescales in our system - the LJ time which is given by 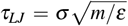^89^ and the Brownian timescale of a free spherical particle, *τ*_*B*_ = *σ* ^2^*/D*, where *D* = *k*_*B*_*T/γ* is the diffusion coefficient and *γ* = 1 is the scaled friction constant. As usual, we set the two timescales equal in our simulation, and hence *τ* = *τ*_*LJ*_ = *τ*_*B*_ = *m/γ*. We also choose the mass of each particle to be unity, i.e. *m* = 1. The temperature of the system is maintained at *T* = *ε/k*_*B*_. We choose a integration time step of *dt* = 0.002*τ*. All results shown in this manuscript are obtained from at least 5 independent simulation runs up to 3 × 10^8^ MD time steps for each set of parameter values. Each simulation is equilibrated for 5 × 10^6^ MD time steps and all quantities are averaged over at least 1000 independent polymer conformations obtained post-equilibration.

## III. RESULTS

First, we characterise how the binder and monomer valencies impact the extent of compaction and reswelling of the chromatin polymer. As the amount of compaction and reswelling differs for different values of BV and MV, we introduce a compaction parameter Ω to denote the extent of compaction/reswelling. The compaction parameter represents the fractional extent of compaction/reswelling compared to the maximum compaction observed for a given set of values of BV and MV and can be expressed as

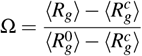

where 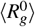 is the average radius of gyration in absence of any binders and 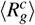 is the lowest average *R*_g_ value (maximum compaction) for a given set of BV and MV values.

Most previous polymer-binder studies, particularly in the context of chromatin, have overlooked the explicit valencies of binders and monomers, often interpreting the polymer-binder interaction strength as a proxy for valency. This interpretation presents two key issues: (a) since the interaction between the polymer and binders is symmetric, it inherently assumes equal valencies for both binders and monomers, and (b) while an increase in interaction strength typically enhances polymer compaction, mirroring the effects of increased valency, it does not necessarily lead to reduced polymer swelling at higher binder concentrations, which would be expected for higher valencies (Fig. S1).

Having established the importance of considering the valencies of binders and monomers explicitly, we now systematically vary these valencies to investigate their individual effects. Fig. 1 shows the variation of *R*_*g*_ and Ω with binder number *N*_*b*_ and binder concentration (*c*), respectively, for different binder valencies (BV) at fixed monomer valencies (MV). Similar to our previous work^83^, the polymer size (*R*_*g*_) initially decreases with increasing *N*_*b*_ and finally starts to increase at higher values of *N*_*b*_ (Fig. 1a-c). Note that for any fixed monomer valency, the polymer size (*R*_*g*_) decreases with increasing binder valency for all binder numbers, as shown in Fig. 1a-c. Increasing BV leads to an effective monomer cloud around binders where each binder binds to multiple monomers which surround the binder itself. This monomer cloud drives the polymer to higher compaction. For low binder valency (BV=2), there is no significant compaction of the polymer irrespective of monomer valency. This is due to the fact that when binders bind to one monomer, it is highly likely for the binder to exhaust its valency by binding to another monomer that is sequentially close along the polymer backbone to the already bound monomer. Hence most of the loops are short-ranged and lack of long-range loops leads to no compaction of the polymer. Although the reswelled *R*_*g*_ of the polymer varies for different values of BV and MV (Fig. 1a-c), interestingly, the fractional reswelling of the polymer (denoted by compaction parameter Ω) for BV ≥ 4 has similar values for a given value of MV as shown in Fig. 1d-f. Therefore, irrespective of the monomer valency, the polymer compaction (and not the reswelling) predominantly depends on the binder valency and the extent of compaction increases with increasing the binder valency. Although this compaction and reswelling is studied for a simple model where all monomers can interact with the binding proteins, in reality, the specific binding sites of these bridging proteins are distributed irregularly on the chromatin chain. We verify that this effect of initial collapse followed by subsequent reswelling is qualitatively the same even for heteropolymer models of chromatin. We find that the qualitative feature of coil-globule-coil transition is also present for two simple heteropolymer models - random heteropolymer and block copolymer (Fig. S2).

**FIG. 1.**
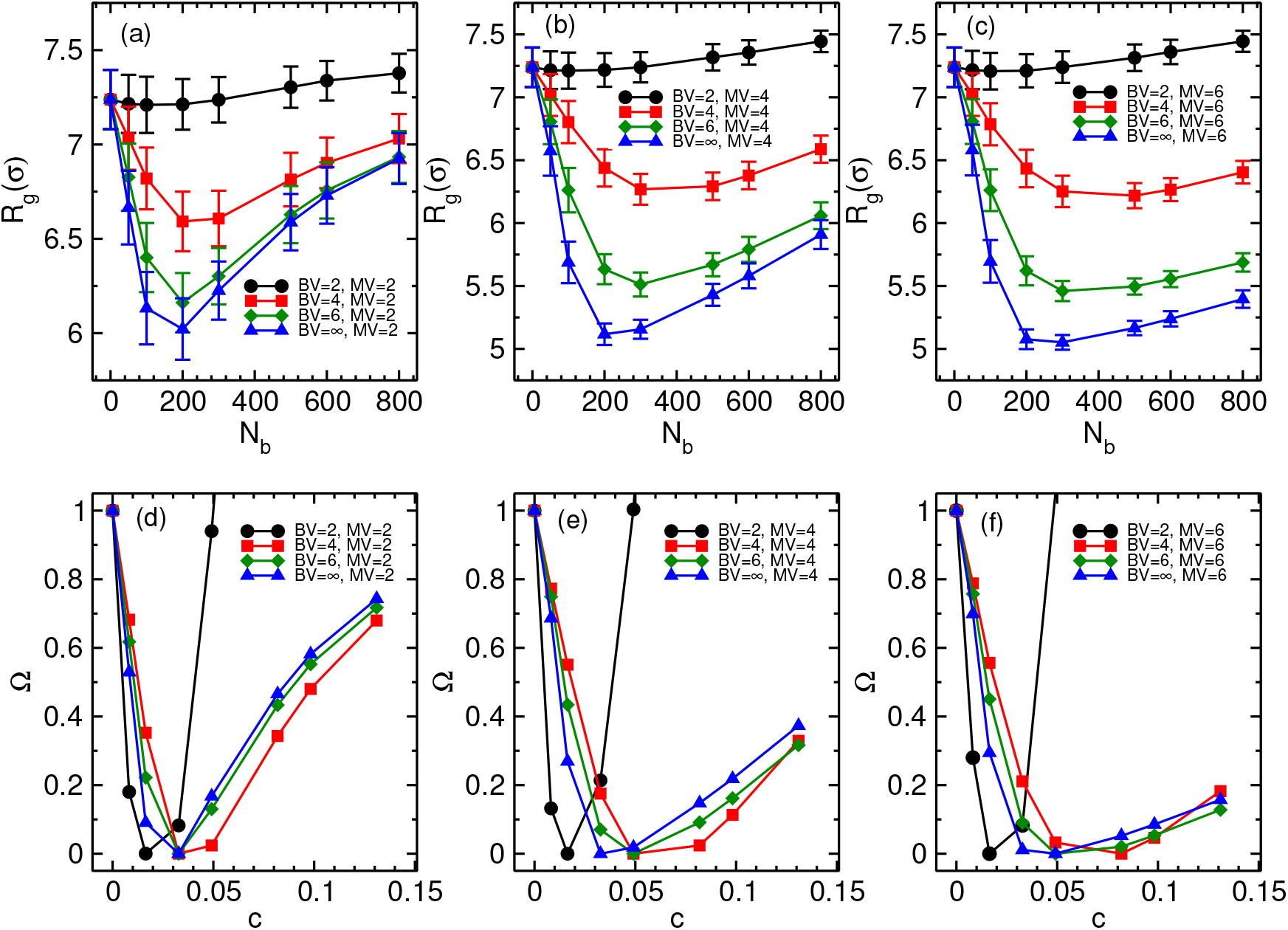
Comparison of variation in *R*_*g*_ as a function of *N*_*b*_ for different binder valencies at fixed monomer valency. Top panels (a-c) shows the variation of *R*_*g*_ as a function of *N*_*b*_ for three different monomer valencies. Each panel compares the the *R*_*g*_ variation for different binder valencies at a fixed monomer valency. The bottom panels (d-f) depict the variation of the compaction parameter (Ω) with binder number density (*c*) corresponding to the same binder and monomer valencies as in the top panels.

To understand the impact of monomer valency on polymer conformation, we now study the variation of *R*_*g*_ and Ω as a function of *N*_*b*_ and *c*, respectively, for different MV values for a fixed BV (Fig. 2). As mentioned earlier, for BV=2, there is a minor compaction of the polymer (Fig. 2a). Surprisingly, the polymer *R*_*g*_, for BV=2, varies identically for MV ≥ 4. For higher binder valencies (BV ≥ 4), the extent of polymer compaction increases as monomer valency is increased, but the reswelling considerably decreases with increasing monomer valencies, as shown in Fig. 2b and 2c. This is also reflected in the corresponding Ω vs *c* plots shown in Fig. 2d-f. Note that the high value of Ω in BV=2 case (Fig. 2d) is an artifact due to the minimal compaction of the polymer. In contrast to the compaction, the extent of reswelling of the polymer predominantly depends on monomer valency as is evident from Fig. 2. Interestingly, the monomer valency has opposite effects on reswelling depending on the value of binder valency. For high binder valency (BV ≥ 4), the value of polymer *R*_*g*_ is lower for higher monomer valencies (Fig. 2b-c). However, for binder valency (BV=2), the *R*_*g*_ of the reswollen polymer is higher for higher monomer valencies (MV ≥ 4) than the reswollen polymer *R*_*g*_ in case of MV=2 (Fig. 2a).

**FIG. 2.**
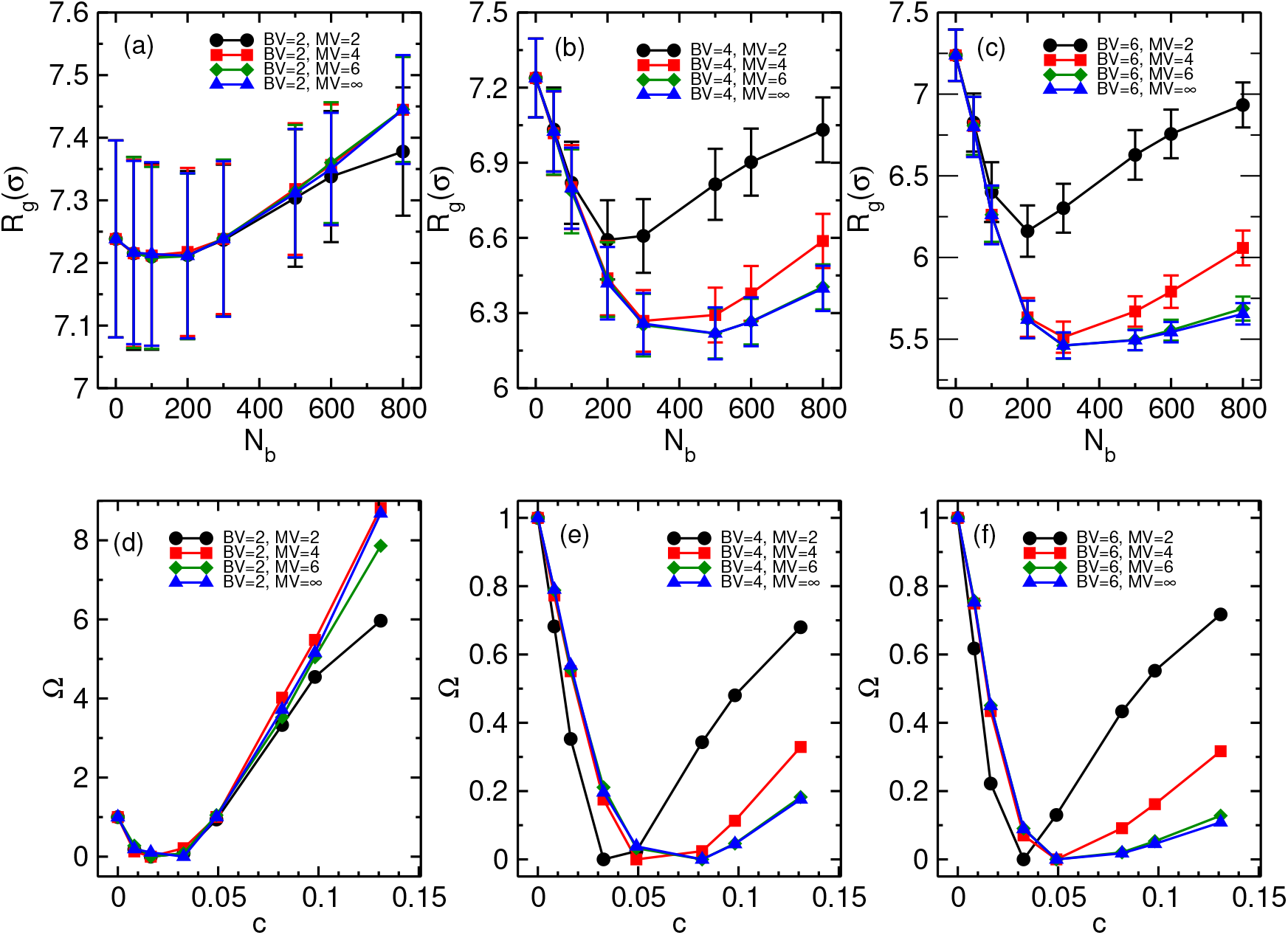
Comparison of variation in *R*_*g*_ as a function of *N*_*b*_ for different monomer valencies at fixed binder valency. Top panels (a-c) shows the variation of *R*_*g*_ as a function of *N*_*b*_ for three different binder valencies. Each panel compares the the *R*_*g*_ variation for different monomer valencies at a fixed binder valency. The bottom panels (d-f) depict the variation of the compaction parameter (Ω) with binder number density (*C*) corresponding to the same binder and monomer valencies as in the top panels.

To understand these opposite effects of monomer valency on polymer reswelling, we now look into the loop size distribution of the system. Specifically, we compare the loop size distributions for the highest number of binders used in our study (*N*_*b*_ = 800) across different binder valency (BV) and monomer valency (MV). We found that for a given monomer valency, increasing binder valency leads to an increase in the number of loops (Fig. 3a-c) and hence the *R*_*g*_ is lower for higher binder valency. A similar increase in loop number is observed with increasing MV values for BV ≥ 4 (Fig. 3e-f). However, when binder valency is low (BV=2), the loop distribution remains relatively unchanged across different monomer valencies (Fig. 3d). The very low number of long-range loops in this case indicates that the reswelling is due to the excluded volume of binders which induces an effective repulsive force between the binders that are bound to the polymer. This can be visualised as an effective binder cloud formed around the monomers due to multiple binders binding to same monomer. Interestingly, while the *R*_*g*_ remains almost identical for higher monomer valencies (MV ≥ 4), the amount of reswelling in case of MV=2 is lower than the reswelling observed in cases of MV ≥ 4 (Fig. 2a). This can be thought of as follows: in case of MV=2, the binder cloud formed by only two binders cannot efficiently exclude the volume, and the volume excluded by two binder clouds can overlap as schematically shown in Fig. 3g. With higher monomer valency, the binder cloud formed by binders around monomers efficiently excludes volumes larger than the individual monomer volume and volumes occupied by the binder clouds cannot overlap (Fig. 3h). This efficient volume exclusion by binder clouds in case of higher monomer valencies increases the effective volume occupied by each monomer and hence reswells the polymer. The more efficient volume exclusion by binder clouds for MV ≥ 4 leads to higher *R*_*g*_ than in the case of MV=2. But with higher binder valency (BV ≥ 4), the number of long loops increases with increasing monomer valency (Fig. 3e-f) and hence the extent of polymer compaction is more for higher monomer valencies (Fig. 2b-c). Due to the increase in the number of long loops with increasing monomer valencies, the extent of reswelling in this case also diminishes.

**FIG. 3.**
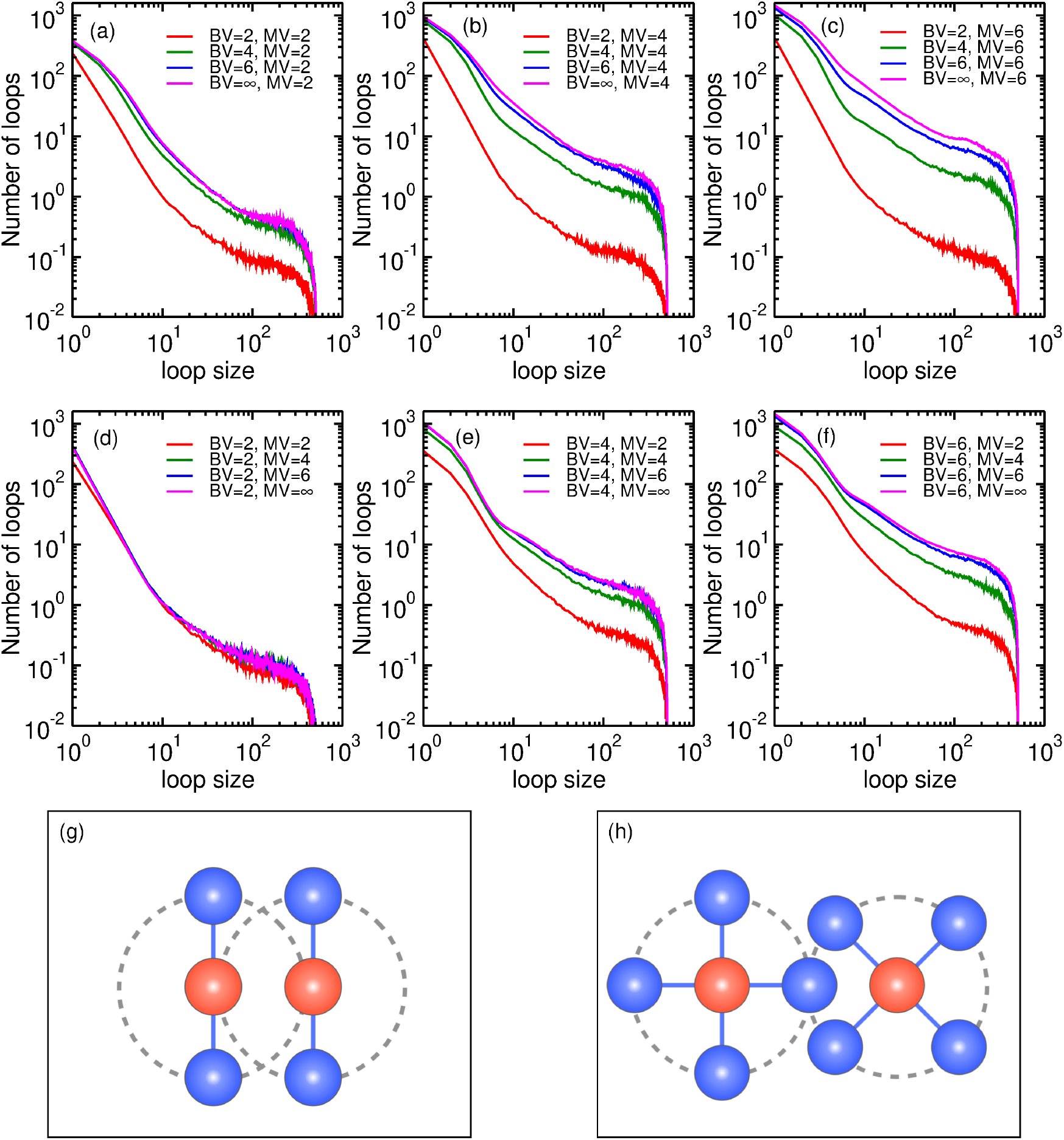
Loop distribution for different binder and monomer valencies for *N*_*b*_ = 800. Panels (a-c) compare the loop size distribution for different binder valencies for 3 fixed monomer valencies, whereas panels (d-f) compare the same for different monomer valencies for 3 fixed binder valencies. Panels (g) and (h) show the excluded volume effects by 2 binders (denoted by blue beads) bound to each monomers and 4 binders bound to each monomers (denoted by red beads), respectively.

Next we plot the loop size distribution for three different binder numbers, each corresponding to a different chromatin conformation - semi-open, fully collapsed, and reswelled (Fig. 4a and 4b). Since the amount of reswell is different for different valencies, we compare the reswelled polymer configurations at same binder number (*N*_*b*_ = 800). Interestingly, for a given binder and monomer valency, two distinct causes of reswelling emerge. One is that reswelling is driven by the loss of long loops at higher *N*_*b*_, while the other is predominantly due to the excluded volume of the binder cloud around each monomer. In the first scenario, when the binder number is low (semi-open polymer configurations), there are fewer long loops. However, as the number of binders increases, leading to a fully collapsed polymer conformation, the number of loops, both long-range and short-range, increases significantly(Fig. 4a). Due to this increase in long-range loops, the polymer collapses and reaches minimum *R*_*g*_. As the binder number increases further, the number of long-range loops diminishes. This can be understood as follows: the compaction of a polymer chain occurs hierarchically across multiple lengthscales^94^. Initially, binders attach to the polymer and induce local compaction, forming small globule-like structures. Subsequently, these local globules merge, driving higher-order compaction and leading to a globally compact globule conformation along the polymer chain. In a valency-restricted scenario at high binder density, binders rapidly form multiple local interactions and saturate the valencies of nearby monomers. While this promotes local compaction at shorter lengthscales, the exhaustion of monomer valencies limits the formation of long-range loops required for global compaction.

**FIG. 4.**
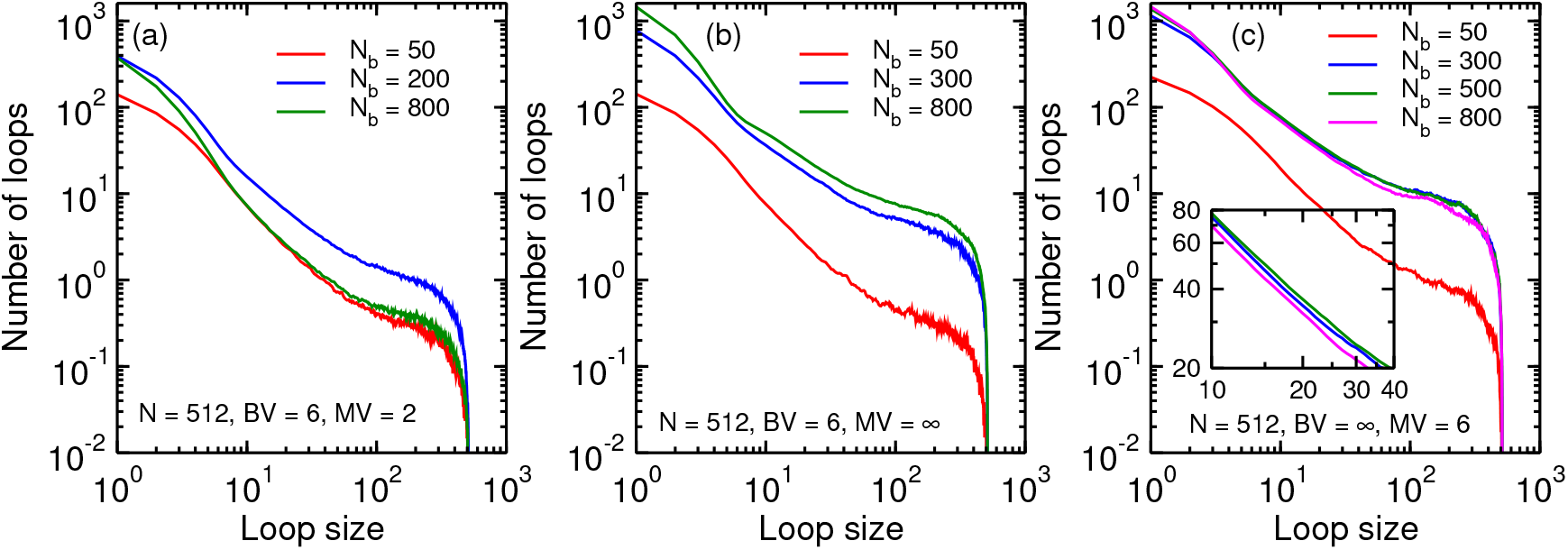
Loop size distribution for different binder and monomer valencies. Panel (a) shows a re-entrant like behaviour in loop size distribution with increasing binder number while panel (b) shows a monotonic increase in number of loops with increasing binder numbers. Panel (c) depicts the transition of excluded volume-driven reswelling into reswelling driven by loss of loops. The inset in panel (c) shows a zoomed-in picture of the loop distribution and clearly show the loss of loops.

This is evident in the significant decrease in long-range loops observed when the binder number is increased from *N*_*b*_ = 200 to *N*_*b*_ = 800, while the number of short-range loops remains nearly unchanged (Fig. 4a). Hence, this reduction of long-range loops at high binder numbers leads to a semi-open state of the polymer which corresponds to higher *R*_*g*_, i.e, the polymer reswells. In contrast, when reswelling of the polymer is dominated by excluded volume effects, the number of long-range as well as short-range loops monotonically increases with increasing binder number (Fig. 4b). This increase in loop number across all loop sizes indicates an increase in the total number of binders within the volume occupied by the polymer. As a result of the mutual repulsion between binders within the polymer’s occupied volume, the polymer undergoes reswelling. The extent of reswelling of chromatin driven by this excluded volume mechanism is strongly influenced by various physical properties, as well as environment of the system. For instance, an increase in the number of binders would mean that more binders are located within the volume occupied by the polymer chain and hence due to their excluded volume the polymer size would increase. Additionally, since larger binders exclude a larger volume, an increase in binder size would result in a more pronounced reswelling (Fig. S3). Furthermore, in our previous study^83^, we have shown that the extent of reswelling also varies depending on the confinement radius, *R*. Interestingly, at even higher binder numbers, reswelling caused by excluded volume effects reaches a saturation point, after which further polymer reswelling is once again dominated by the loss of loops. As illustrated in Fig. 4c, when the binder number is increased from *N*_*b*_ = 300 (corresponding to the fully compacted conformation) to *N*_*b*_ = 500, the number of loops increases, suggesting that reswelling in this case is primarily driven by excluded volume effects. However, as the binder number increases further to *N*_*b*_ = 800, the number of long-range loops decreases, as shown in the inset of Fig. 4c. Consequently, the additional reswelling of the polymer from *N*_*b*_ = 500 to *N*_*b*_ = 800 is clearly dominated by the loss of loops.

Now that we have identified two distinct physical mechanisms responsible for polymer reswelling, we plot a classification map to distinguish the dominant mechanism of reswelling for different combinations of binder and monomer valencies (Fig. 5). It should be noted that the two mechanisms are not mutually exclusive and both contribute simultaneously in the polymer reswelling as previously described in Fig. 4c. We thus define a numerical threshold to distinguish the regimes depending on the dominant cause of the reswelling for a given set of binder and monomer valencies. As at very high binder numbers all reswelling phenomena are expected to be dominated by loss of loops, we restrict our classification map to the binder number *N*_*b*_ = 800 (equivalent to binder volume density *φ*_*b*_ ∼ 0.7 and binder number density *c ∼* 0.13) (Fig. S4). Therefore, in the presence of 800 binders, if the number of loops is lower than the number of loops at lower binder numbers, *N*_*b*_ = 600 (*φ*_*b*_ ∼ 0.5 and *c* ∼ 0.13), then we call it a loop loss regime, and if the number of loops monotonically increases till *N*_*b*_ = 800 then we call it excluded volume regime. As discussed earlier, for BV=2, the number of long loops is low and hence, as expected, we found that the reswelling in this case is dominated by excluded volume effects. We also found that reswelling dominated by the loss of loops is prevalent in lower monomer valencies, while at higher monomer valencies, reswelling is predominantly driven by excluded volume effects.

**FIG. 5.**
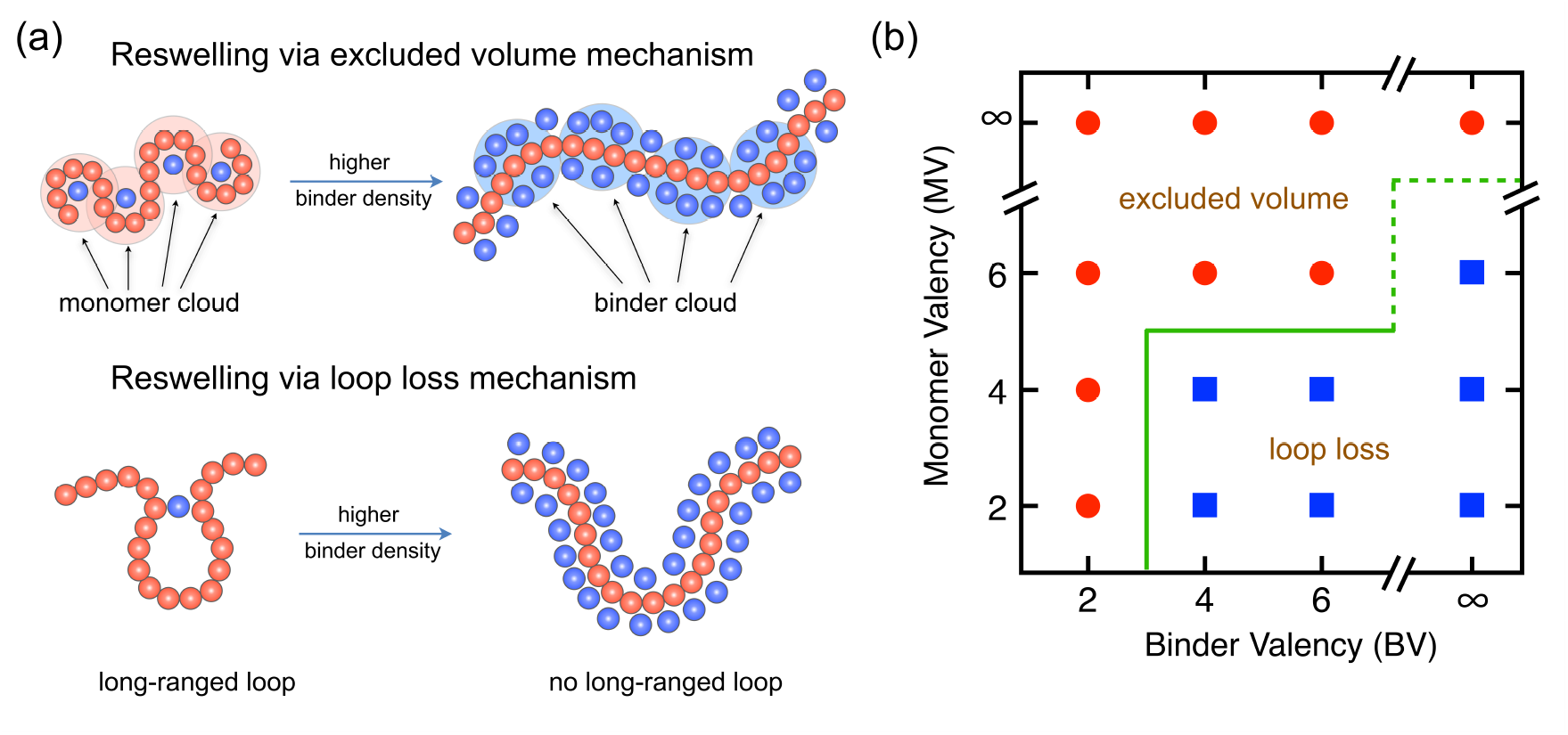
Distinct mechanisms of polymer reswelling. Panel (a) shows schematic representation of two mechanisms of polymer reswelling. Red chain denotes the polymer and the blue beads depict the binders. Panel (b) shows a classification map distinguishing the two regimes of reswelling mechanism on binder valency-monomer valency plane. Circles (red) denote reswelling due to excluded volume while squares (blue) denote reswelling due to loss of long-range loops. The green line denotes the boundary between the two mechanisms.

## IV. DISCUSSION

Several polymer models have been studied in the literature in the context of chromatin organisation. While chromatin compaction is well understood using polymer folding in presence of multivalent bridging molecules, very little is known regarding the role of these bridging proteins in reswelling the chromatin. In our previous work^83^, we have shown that these multivalent binders can both compact and reswell the chromatin polymer under confinement. Also, a chromatin monomer can have multiple binding sites for these proteins, making the polymer multivalent. Therefore, it is important to understand how the valencies of binders and monomers affect the compaction and reswelling of the chromatin under confinement.

In this work, we have shown that the binder and monomer valencies can play a crucial role in chromatin organisation inside cell nucleus. We have found that the extent of chromatin compaction increases with increasing binder valency. Additionally, we do not observe any significant compaction of the polymer in the case of binder valency 2, as in this case most loops are short-ranged and lack of long-range loops results in no compaction. The polymer still shows a reswelling at higher binder numbers for binder valency 2. This reswelling of polymer is found to be less for monomer valency 2, while for higher monomer valencies the reswelling is identical and higher than the reswelling for monomer valency 2. On the other hand, the extent of reswelling is found to diminish with the increasing monomer valency for higher binder valencies. We then further quantified the loop size distributions to understand this contrasting behaviour of polymer size for different monomer valencies. We show that in one case the number of long-range loops monotonically increase with increasing binder number, while in other case the loop size distributions show a non-monotonic behaviour with increasing binder numbers. We conclude that the reswelling of the polymer is driven by two factors - excluded volume effects of the binders and lack of long-range loops at higher binder numbers. While the monotonic increase in number of loops contributes to the increasing excluded volume effects and the non-monotonic variation in the loop distribution indicates a loss of loops at higher binder numbers. Based on these features we obtain a classification map separating two causes of reswelling as a function of monomer and binder valencies. We have found that at higher binder valency and lower monomer valency regimes the reswelling is predominantly driven by loss of long-range loops, while in the other regimes reswelling is driven by excluded volume effects. Finally, we have also shown that at even higher binder numbers, excluded volume driven reswelling saturates, and further reswelling with increasing binder number is dominated by loss of long-range loops.

These differential causes of reswelling for different binder and monomer valencies can have important biological implications. Different experimental studies have reported that various DNA-associated proteins have different sizes ranging from 10*nm* to a few hundred *nm*^49,95^. Furthermore, transcription factors are known to aggregate near their binding sites and form large transcriptional condensates^96–98^. In case of excluded volume-dominated reswelling, the size of the binder proteins or transcriptional condensates would play a crucial role. Larger proteins or condensates exclude a larger volume, and thus the polymer reswelling would be larger, while a smaller protein or condensate would exclude less volume, and the reswelling would not be as significant as in case of bigger proteins. However, when reswelling is dominated by loss of loops, the extent of reswelling is independent of protein size. Certainly, future experiments that specifically focus on the number of loops can provide valuable insights into the underlying physical mechanisms responsible for chromatin reswelling in the presence of particular DNA-associated proteins. By systematically varying the binder and monomer valencies and observing the resulting changes in chromatin compaction and reswelling, researchers can gain a deeper understanding of the specific factors driving these processes. This knowledge can help elucidate the roles of different DNA-associated proteins and their interactions with chromatin in the stability and dynamics of chromatin organisation.

In summary, our simple polymer-binder system in confinement has been able to illustrate the counter-intuitive effects of binder and monomer valencies on polymer conformations. In addition, we found two distinct mechanisms that are responsible for polymer swelling - *excluded volume effects* and *loss of long-range loops*. The extent of collapse and reswelling will also depend on the distribution of specific binding sites along chromatin, and future work with a systematic study of real chromatin sequences with specific binding sites can help uncover sequence-specific modulations of this phenomenon. Further, we have considered that proteins only interact via excluded volume effects. In vivo, multiple proteins are known to undergo oligomerisation and therefore allowing proteinprotein interactions together with multivalency can yield interesting insights into chromatin organisation^60,99,100^. Despite these assumptions, our simple homopolymer model captures the basic underlying physical mechanism of chromatin organisation and the effects of proteins and chromatin valencies.

## SUPPLEMENTARY INFORMATION

The supplementary information contains additional data.

## ACKNOWLEDGEMENTS

The author would like to thank Mithun K. Mitra and Shuvadip Dutta for their valuable discussions and detailed feed-back on the manuscript. The author also acknowledges the computing facilities and partial financial support provided by IIT Bombay. Additionally, The author would like to express his gratitude to INFN Napoli for their financial support through the “Postdoctoral Senior Level 3 Research Grant in Theoretical Physics” (24736/2022).

## DATA AVAILABILITY STATEMENT

Data supporting the findings of this study are available from the corresponding author on reasonable request.

