## Supplementary File for "Binder and monomer valencies determine the extent of collapse and reswelling of chromatin"

Sougata Guha[1,2]

[1]*Department of Physics, IIT Bombay, Mumbai 400076, India*
[2]*INFN Napoli, Complesso Universitario di Monte S. Angelo, 80126 Napoli, Italy*
**

### Supplementary Information

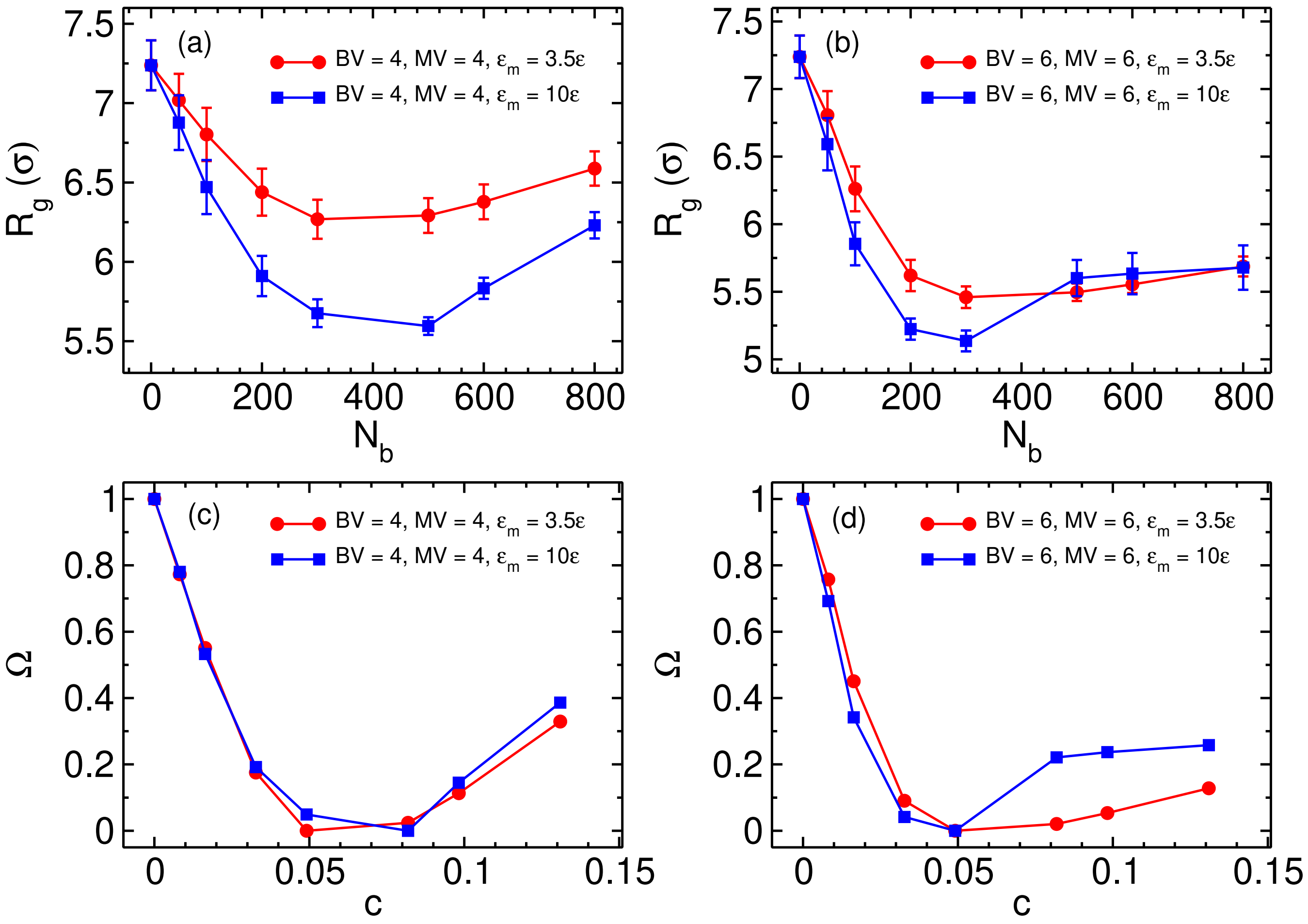


Fig. S1. Comparison of polymer compaction and reswelling for two different interaction strengths ($\epsilon_m$) in the the cases of BV=4, MV=4 and BV=6, MV=6.

.

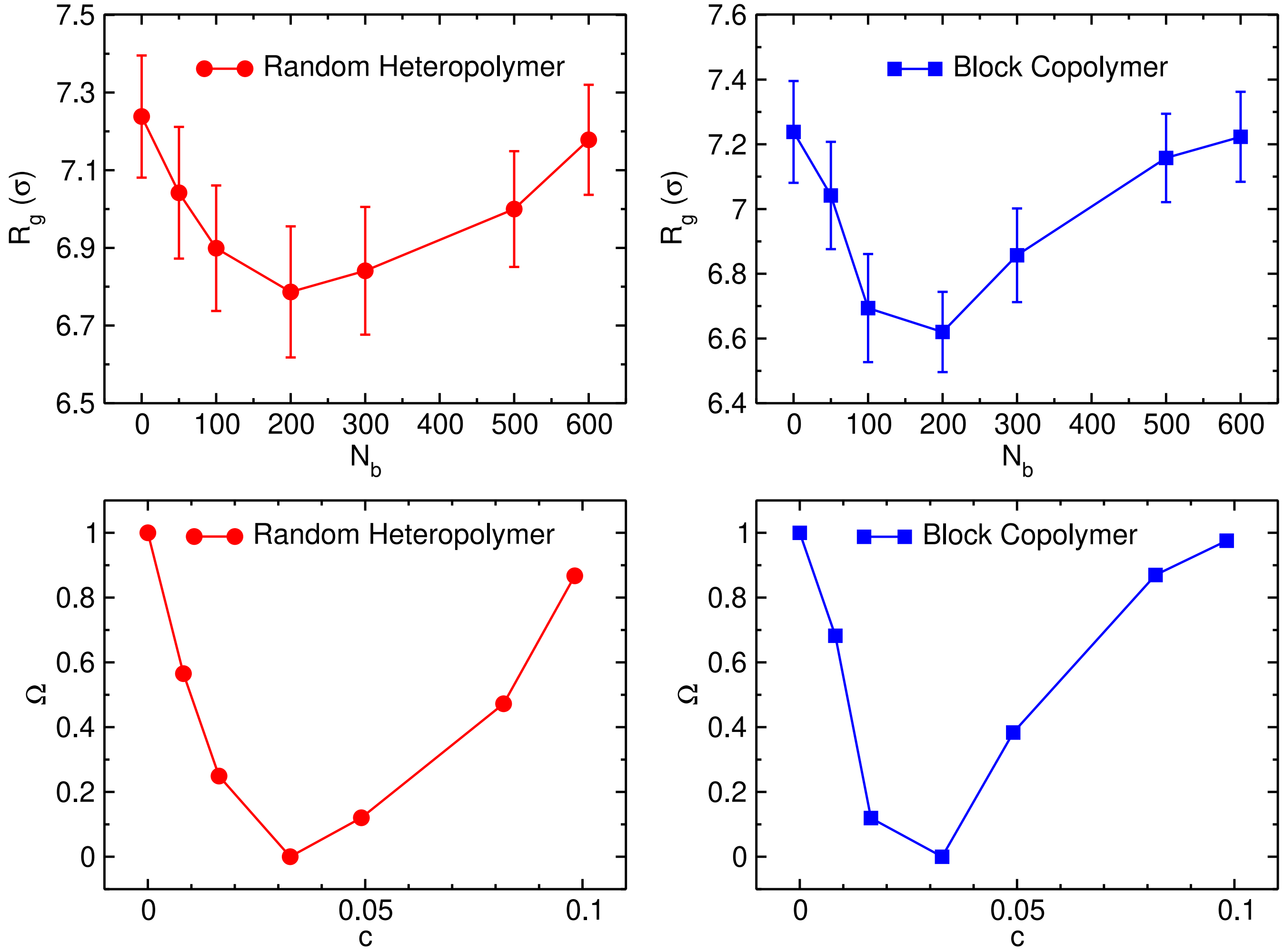


Fig. S2. Variation of $R_g$ and $\Omega$ as functions of $N_b$ and $c$, respectively, for two types of heteropolymers. In both cases, approximately 33% of the monomers (active monomers) are designated as binding sites for the binders, while the remaining monomers (inactive monomers) interact with binders solely via excluded volume interactions. For the random heteropolymer, the active binding sites are distributed randomly along the polymer chain. In contrast, for the block copolymer, an alternating structure is employed, consisting of 10 consecutive active monomers followed by 20 inactive monomers. The binder and monomer valencies in this case are set as BV=4 and MV=4.

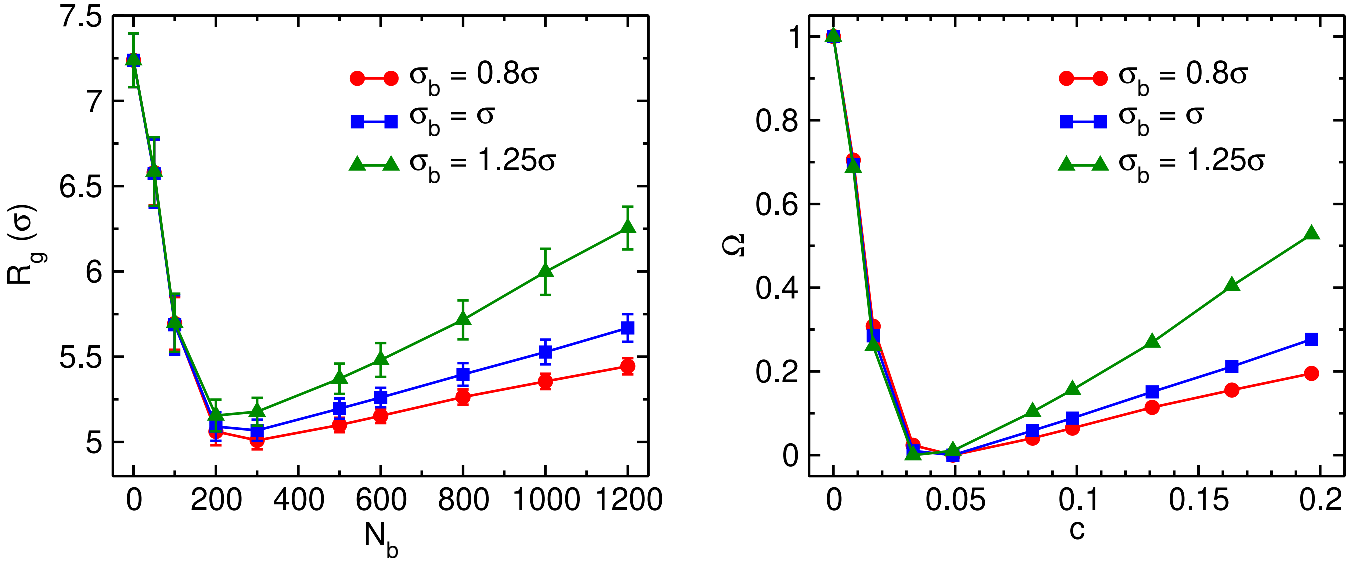


Fig. S3. Comparison between variation of $R_g$ and $\Omega$ as a function of $N_b$ and $c$ respectively for three different binder sizes in the case of BV= $\infty$, MV= $\infty$. The reswelling is higher for larger binder due to larger excluded volume.

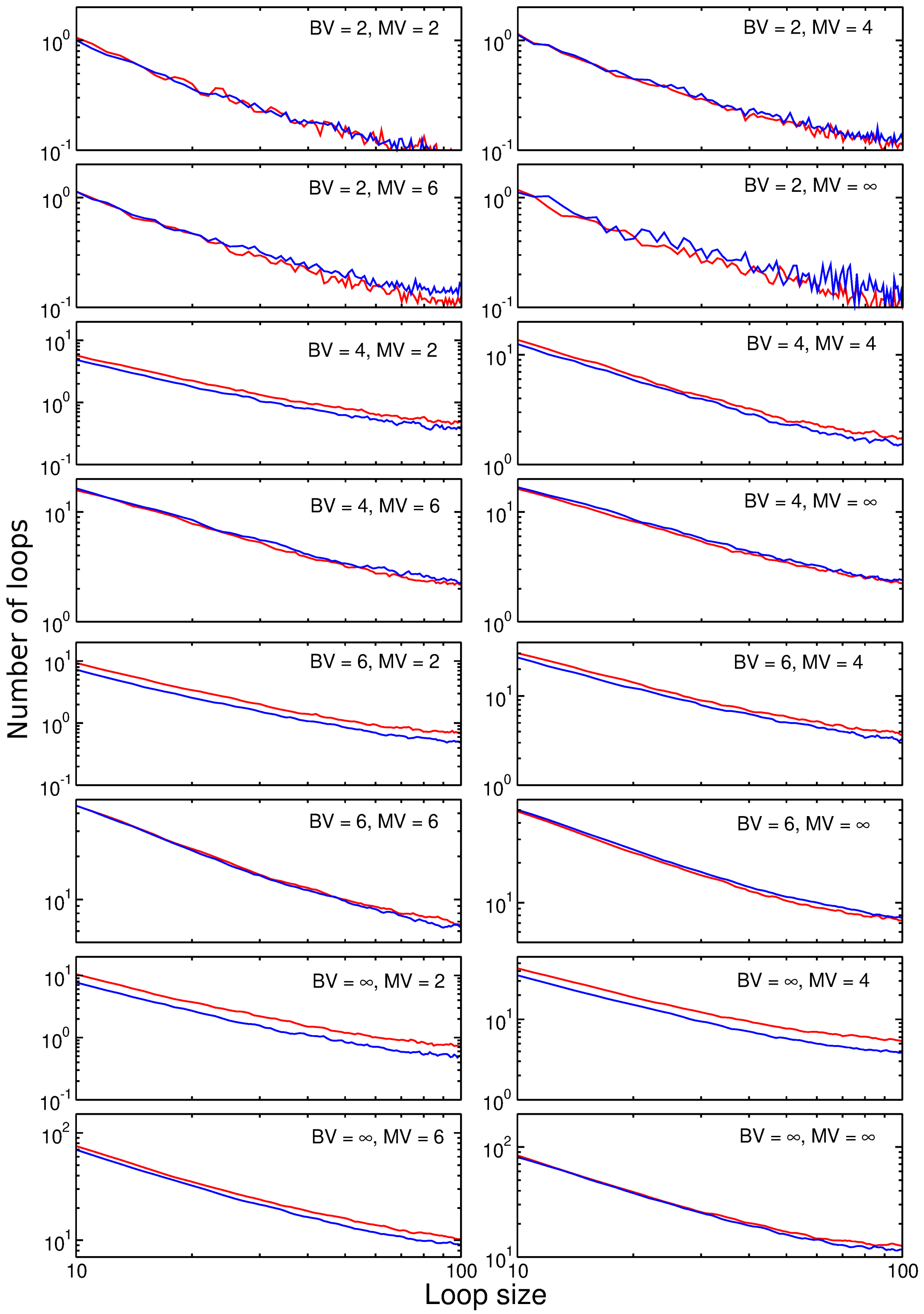


Fig. S4. The loop size distribution for $N_b = 600$ and $N_b = 800$ for different values of BV and MV. The red line corresponds to $N_b = 600$ and the blue line corresponds to $N_b = 800$.
